# Airway-associated macrophages in homeostasis and repair

**DOI:** 10.1101/2020.06.16.146035

**Authors:** Anna E. Engler, Alexandra B. Ysasi, Riley M.F. Pihl, Carlos Villacorta-Martin, Hailey M. Heston, Hanne M.K. Richardson, Noah R. Moniz, Anna C. Belkina, Sarah A. Mazzilli, Jason R. Rock

## Abstract

There is an increasing appreciation for the heterogeneity of myeloid lineages in the respiratory system, but whether distinct populations associate with the conducting airways remains unknown. We use single cell RNA sequencing, flow cytometry and immunofluorescence to characterize myeloid cells of the mouse trachea during homeostasis and epithelial injury/repair. We identify submucosal macrophages that are similar to lung interstitial macrophages and intraepithelial macrophages, and find that repair of the tracheal epithelium is impaired in *Ccr2*-deficient mice. Following injury there are early increases in neutrophils and submucosal macrophages, including M2-like macrophages. Unexpectedly, intraepithelial macrophages are initially lost but later replaced from CCR2^+^ monocytes. Mast cells and group 2 innate lymphoid cells are sources of IL13 that polarizes macrophages and directly influences basal cell behaviors. Their proximity to the airway epithelium establishes these myeloid populations as potential therapeutic targets for airway disease.

## Introduction

The respiratory system can be divided into at least two functional zones – the proximal conducting airways and the distal alveoli – and each contains distinct populations of epithelial and mesenchymal cells with unique functions (Rock and Hogan, 2011). Alveolar macrophages (AMs) are thought to be the tissue resident macrophages of the lung, and there is evidence for ontological, molecular and functional heterogeneity in this population (Misharin et al., 2013; Tan and Krasnow, 2016). Much less is known about the myeloid lineages of the airways, but recent evidence points to molecularly distinct interstitial macrophages (IMs) associated with the submucosa (Gibbings et al., 2017). We have exploited the mouse trachea as a model for smaller airways, including those of humans (Rock et al., 2010) to learn more about these cells during homeostasis and repair after injury.

Recent studies have demonstrated roles for immune modulation of lung repair and regeneration. We recently demonstrated that CCR2^+^ inflammatory monocytes and M2-like Arg1^+^ macrophages promote regeneration from type 2 alveolar epithelial stem cells in a mouse model of partial pneumonectomy (Lechner et al., 2017). In the airways, it is well known that Th2 cytokines influence the cellular composition of the epithelium (Danahay et al., 2015; Kuperman et al., 2002), but less is known about how myeloid lineages impact injury and repair. To address this question, we used flow cytometry and single cell RNA sequencing (SCSeq) to characterize the myeloid populations in the mouse trachea at steady state and during repair of the epithelium following chemically-induced injury, when the number of macrophages increases (Tadokoro et al., 2014). We provide evidence for airway-associated macrophages that are distinct from AMs and show that their polarization shifts from M0-like to M2-like during epithelial repair. These M2-like macrophages originate, at least in part, from a *Ccr2^+^* precursor and regeneration of the tracheal epithelium is impaired in the absence of *Ccr2* expression. Lastly, we identify a unique macrophage subset that is located in close proximity to airway basal cells within the tracheal epithelium. Our data suggest that these cells are lost along with the tracheal epithelium after chemically-induced injury and are replaced by CCR2^+^ cells during epithelial repair.

## Results

### The trachea contains interstitial macrophages under homeostatic conditions

The mouse trachea provides a well-characterized model of the pseudostratified airway epithelium. To characterize the macrophages associated with this tissue we first performed flow cytometry on cells from the dissociated normal trachea and from the remaining lung lobes for comparison. This identified a population of CD45^+^;F4/80^+^ myeloid cells in the trachea which are negative for SiglecF (Figure S1A-B), a classical marker of alveolar macrophages (AMs). Approximately 38% of the CD45^+^ cells in the trachea are MerTK^+^. Most of these are CD11b^hi^ and variably express CD11c (FigureS1B), a phenotype consistent with interstitial macrophages (IM) previously described for the distal lung. In contrast, the majority of CD45^+^;MerTK^+^ cells in the lung are Cd11b^neg/lo^;CD11c^hi^ AMs with a minor population of MerTK^+^;CD11b^hi^ IMs (Figure S1B and (Gibbings et al., 2017)).

### Tracheal macrophage dynamics during injury and repair of the tracheal epithelium

Following infection, IMs expand through local proliferation and modulate lung inflammation (Kawano et al., 2016; Ural et al., 2020). We used a model of polidocanol-induced injury to determine how the tracheal macrophage populations respond to non-infectious injury of the airway epithelium (Borthwick et al., 2001). We analyzed the tracheas of polidocanol injured and PBS treated sham control animals 1-, 3- and 7-days post injury (hereafter 1dpP, 3dpP and 7dpP) (Figure S1C-D). Polidocanol results in the death and sloughing of suprabasal ciliated and secretory cells of the pseudostratified epithelium within 3 hours, largely sparing basal cells (Figure S1D). The normal proportions of epithelial cells and epithelial morphology are restored within 1-2 weeks (Figure S1D) by the proliferation and differentiation of basal cells (Borthwick et al., 2001; Plasschaert et al., 2018)). At 1 dpP we observed an increase in the number of CD45^+^ cells in the mesenchyme beneath the injured epithelium (Figure 1A). Flow cytometry showed a rapid increase of CD45^+^ cells and CD45^+^F4/80^+^ cells within 1dpP (Figure 1B). We then carried out a more detailed analysis using a 15-color myeloid immunophenotyping panel (Supplemental Table S1). Opt-SNE dimensionality reduction and PhenoGraph unsupervised clustering of live, extravascular CD45^+^ cells from sham and 1, 3 and 7 dpP tracheas showed 24 distinct clusters (Figure 1C, S1E), of which 17 were categorized as myeloid populations (Figure 1D). The proportion of cells expressing a cell surface phenotype consistent with neutrophils rapidly increased 1 dpP compared to controls (Fig 1C and E, Clusters 10, 2 and 3). We also observed increases in the numbers of Ly6C^+^ putative newly recruited macrophages (Cluster 9) and tissue macrophages (Cluster 4) at 3 dpP. Within 7 dpP, cellular composition had returned to baseline (Figure 1E-F). These data reveal dynamic changes in the myeloid populations during repair and implicate monocyte recruitment as a potential contributing mechanism.

**Figure 1:**
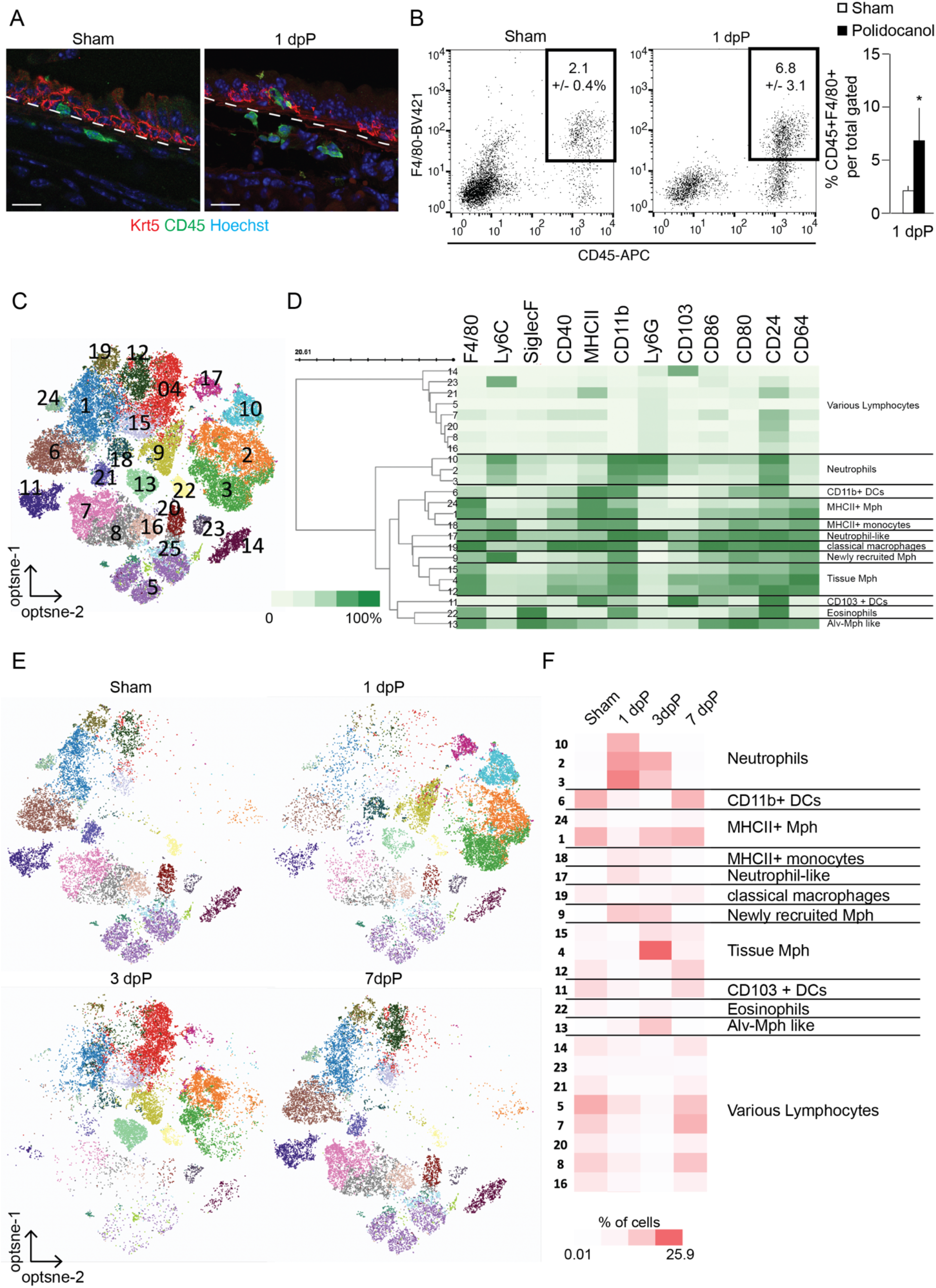
Temporal changes of the tracheal myeloid compartment following injury; (A) Immunofluorescence on sections of trachea show an increase in the number of CD45^+^ cells (green) near basal cells (KRT5, red) following polidocanol-induced injury. Scale bar 25 μm (B) FACS analysis of CD45^+^;F4/80^+^ myeloid cells after 1dpP; Percentage of CD45^+^F4/80^+^ cells of total (n=4); (C) Unsupervised clustering of all cells combined from sham, 1-, 3-, and 7dpP based on 15-marker myeloid panel; (n=1/timepoint, consistent of 2 animals) (D) Heatmap of marker expression data upon which cluster annotations were based, scaled from 0 to 100% of normalized, arcsin value of fluorescent intensity (E) Cells isolated on indicated days post-injury, visualized by cluster assignment; (F) Relative abundance of cells from 24 distinct clusters across injury and repair, scaled 0.01 to 25.9% of total cells in analysis.

### Transcriptional changes during injury and repair

To further characterize the immune composition during tracheal injury and repair, we performed SCSeq of CD45^+^ cells from dissociated tracheas of uninjured control and mice 1, 4 and 7 dpP. We performed post-processing filtering to focus our analyses on myeloid cells (F4/80^+^, *Adgre1^+^*) (Figure 2A, S2A). Uniform manifold approximation and projection (UMAP) visualization (Figure 2B) and analysis of differential gene expression (Figure S2B, Supplemental Table S2) shows that the myeloid cells from uninjured tracheas and 4 and 7 dpP are largely overlapping and these are enriched for classic macrophage and monocyte signatures (Figure 2 C-D). There were also clusters of cells with signatures of dendritic cells (DCs) (*Cd103^+^; Cd11b^+^*), presumably expressing low levels of *Adgre1* detectable by sequencing but not FACS. Upon injury, the number of cells expressing transcripts associated with M2-like macrophages, such as *Arg1* and *Arg2* are increased (Figure 2C, Figure S2B, Supplemental Table S2). Concomitantly the expression of transcripts associated with *Ccr2^+^* and *Cx3cr1^+^* monocytes is rapidly reduced, but returns to baseline 4 dpP and 7 dpP (Figure 2D). We did not observe significant changes in transcripts associated with cell cycle during the course of injury and repair (Figure 2E, gene set Supplemental Table S4).

**Figure 2:**
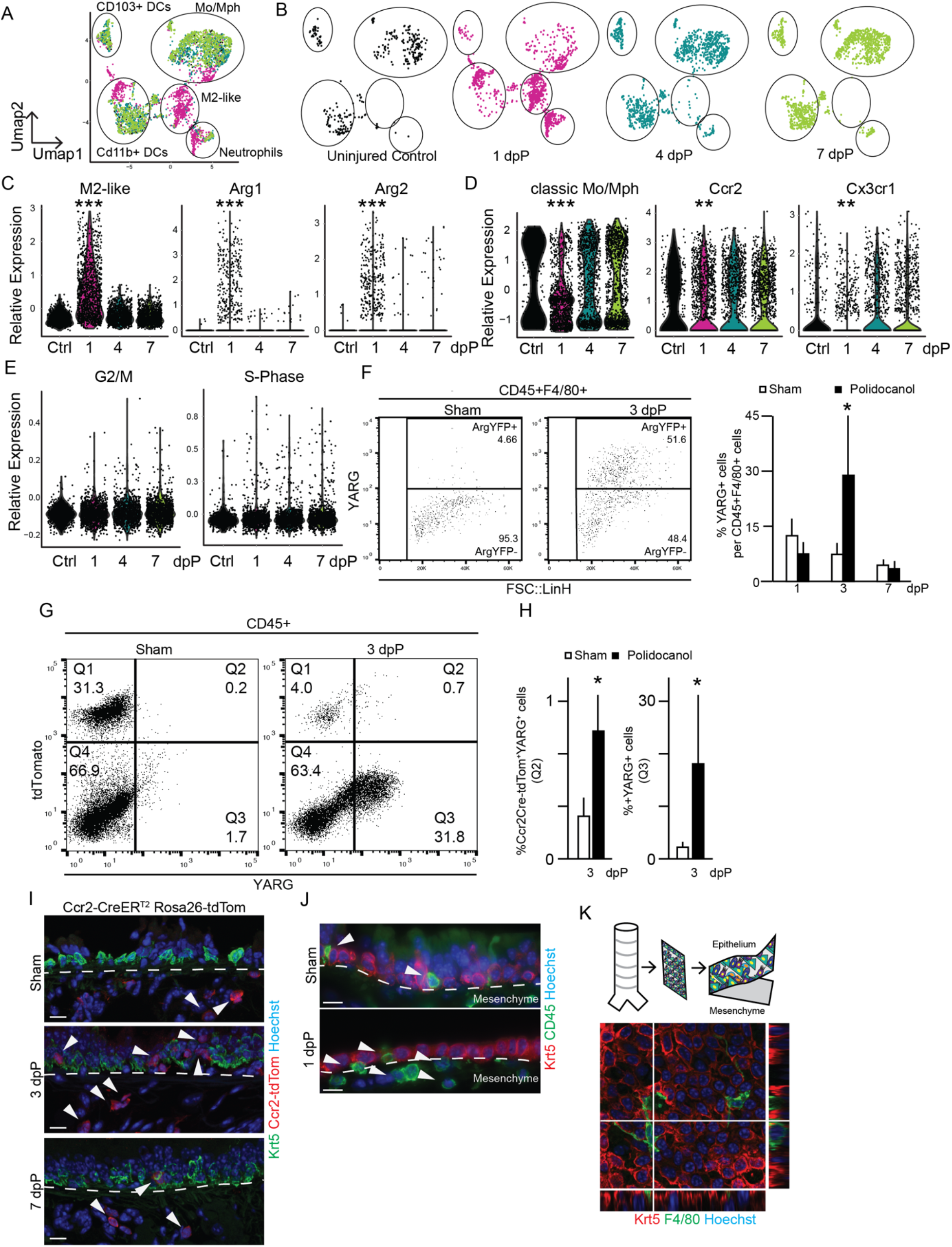
Unbiased analysis of tracheal myeloid populations after injury; (A) UMAP visualization of all SCSeq data across airway injury and repair overlaid; (n=1/time point, consistent of 3 animals); captured cells: uninjured=458, 1dpP=1625, 4dpP=1507, 7dpP=1717 (B) UMAP visualization of SCSeq data from individual time points. From left to right, uninjured controls, 1dpP, 4dpP and 7dpP; (C) Violin plots M2-like macrophage gene signature and *Arg1* and *Arg2* (D), Violin plots classic monocyte/macrophage gene signature and *Ccr2* and *Cx3cr1;* (E) FACS Analysis of dissociated tracheas from YARG^+^ mice gated on CD45^+^;F4/80^+^ (n(Sham)=3, n(3dpP)=3); (F) Quantification of YARG FACS data (G) FACS Analysis of dissociated tracheas from Ccr2-CreERT2;ROSA-tdTom, YARG animals after tamoxifen administration and Sham or Polidocanol treatment (n(Sham)=4, n(3dpP)=3); (H) Quantification of FACS results; (I) Immunofluorescence analysis of Ccr2-CreERT2;ROSA-tdTom animals during injury and repair; Krt5 (green), Ccr2-CreER^T2^;ROSA-tdTom lineage trace (red), and nuclear stain Höechst (blue). scale bar 15 μm; (J) Histology of wild type animals; Krt5 (red), CD45 (green), nuclear stain Höechst, scale bar 15 μm; (K) Immunofluorescence on peeled epithelial sheet showing F4/80^+^ myeloid cells (green) Krt5^+^ epithelial basal cells (red). nuclear stain Höechst (blue);

### The number of M2-like macrophages increases following polidocanol-induced injury

We performed flow cytometry to confirm our observations on the protein level. Consistent with earlier data (Figure 1), the number of CD45^+^;F4/80^+^ myeloid cells increased at 1 and 3 dpP and returned to baseline by 7dpP (Figure S2C). We used an Arginase-YFP reporter mouse (YARG, (Reese et al., 2007)) to quantify M2-like macrophages in the trachea by flow cytometry. YARG^+^ cells were uniquely increased at 3dpP compared to sham and other intervals post-injury (Figure 2F, S2C). This translational readout of arginase expression increased slightly later than the M2-like macrophage associated transcripts (Figure 2B).

Because flow cytometry demonstrated an increase in newly recruited monocytes and macrophages 1-3 dpP (Figure 1C-F), we asked whether the M2-like macrophages were derived from a CCR2^+^ monocyte precursor. We administered tamoxifen to Ccr2-CreER^T2^;ROSA-tdTom;YARG mice to lineage trace CCR2^+^ monocytes and determine whether they gave rise to YARG^+^ cells (Figure S2D). Two weeks later we injured the tracheal epithelium with polidocanol and analyzed tracheas at 3dpP (Figure S2D). As above, the number of CD45^+^ cells was increased 3dpP compared to sham. At this time the percentage of CD45+ cells that are Tom^+^ cells decreased (Figure 2G), but the total number of Tom+ cells was unchanged (Sham: 3018 +/- 1549; Polidocanol: 2717 +/- 706;). This result presumably reflects the local proliferation and/or recruitment of CCR2-negative leukocytes, including neutrophils, lymphocytes and other monocyte subpopulations (Figure 1 and (Joshi et al., 2020; Ural et al., 2020)). Consistent with our transcription and flow cytometry data, both the total number and proportion of YARG^+^ cells were increased 3dpP compared to controls. There was an approximately 3-fold increase in the proportion of lineage traced, Tom^+^YARG^+^ cells (0.7%), suggesting that CCR2^+^ monocytes can give rise to M2-like macrophages. However, the majority of YARG^+^ cells was not lineage traced, presumably derived from resident macrophages, CCR2^neg^ monocytes or escaped recombination (Figure 2G-H).

We sought to localize cells derived from lineage traced CCR2^+^ monocytes during injury and repair of the tracheal epithelium. We delivered a single dose of tamoxifen to Ccr2-CreER^T2^;ROSA-tdTom mice and injured the epithelium with polidocanol 4 days later. Tracheas were harvest for histology 1, 3 and 7 dpP (Figure S2E). As expected, we observed Tom^+^ cells in the submucosa beneath the tracheal epithelium in sham and injured tracheas at all times examined. To our surprise, we also observed an increased number of Tom^+^ cells within the regenerating epithelial layer of the trachea at 3 and 7 dpP (Figure 2I). This unexpected finding led us to confirm the existence of CD45^+^ leukocytes within the pseudostratified tracheal epithelium of sham and injured wild type mice (Figure 2J). Significantly, very few of these intraepithelial leukocytes are lineage traced in sham Ccr2-CreER^T2^;ROSA-tdTom^+^ mice at any time (Figure 2I). Together these data show: (1) a dramatic increase in the number of YARG^+^ M2-like macrophages after injury, (2) a population of myeloid cells within the tracheal epithelium that is not labelled in Ccr2-CreER^T2^;ROSA-tdTom^+^ mice under steady state conditions and (3) increases in the numbers of Ccr2-CreER^T2^;ROSA-tdTom^+^ lineage traced cells in the tracheal mesenchyme and within the epithelium following injury.

### A population of myeloid cells is found within the pseudostratified tracheal epithelium

Next, we focused on the population of CD45^+^ leukocytes within the tracheal epithelium. We peeled sheets of epithelium away from the underlying mesenchyme following a gentle enzymatic digestion as previously described (Rock et al., 2009). Immunofluorescence on isolated sheets revealed CD45^+^ (Figure S2F) and F4/80^+^ (Figure 2K) myeloid cells within the epithelial layer in close proximity to KRT5^+^ basal cells. We similarly analyzed the tracheas of transgenic mice that express GFP under the control of the CSF1R locus, a broad marker of myeloid lineages (cFMS::eGFP, Figure S2I) (Sasmono et al., 2003). Immunofluorescence shows that the intraepithelial CD45^+^ and F4/80^+^ cells also express cFMS::eGFP^+^ (Figure S2J). We called these cells intraepithelial airway macrophages (IAMs).

### Most intraepithelial myeloid cells are molecularly distinct from other pulmonary myeloid lineages

We used the 15-color myeloid panel (Figure 1, Supplemental Table S1) to compare IAMs to cells from the tracheal mesenchyme and lung parenchyma (containing alveoli and intralobar airways). We performed opt-SNE embedding of flow cytometry data from equal numbers of live, extravascular CD45^+^ cells from each of the 3 tissues (Figure 3A). Cell identities were assigned (Figure 3B) based on marker expression (Supplemental Table S1 and (Misharin et al., 2013)). Because the gating strategy was developed for dissociated lung parenchyma, the majority of cells from this tissue were contained within the conventional gates (Figure 3B). In contrast, the majority of myeloid cells from the tracheal epithelium fell outside of these gates, suggesting novel myeloid populations or cell states associate with the trachea under homeostasis (Figure S3A-B and light grey cells in Figure 3B). The few cells from the tracheal epithelium that fell within traditional gates were mostly macrophages and monocytes (Figure 3B) and a few DCs. The tracheal mesenchyme and lung parenchyma contained abundant CD103^+^ and CD11b^+^ DCs. It is interesting to note that some cell types, such as macrophages and DCs, were observed in all 3 tissues, but exhibited slight differences in cell surface profiles represented as small shifts in tSNE space (Figure 3B). As expected, both the peeled tracheal epithelium and the underlying tracheal mesenchyme were devoid of SiglecF^+^ AMs (Figure 3B).

**Figure 3:**
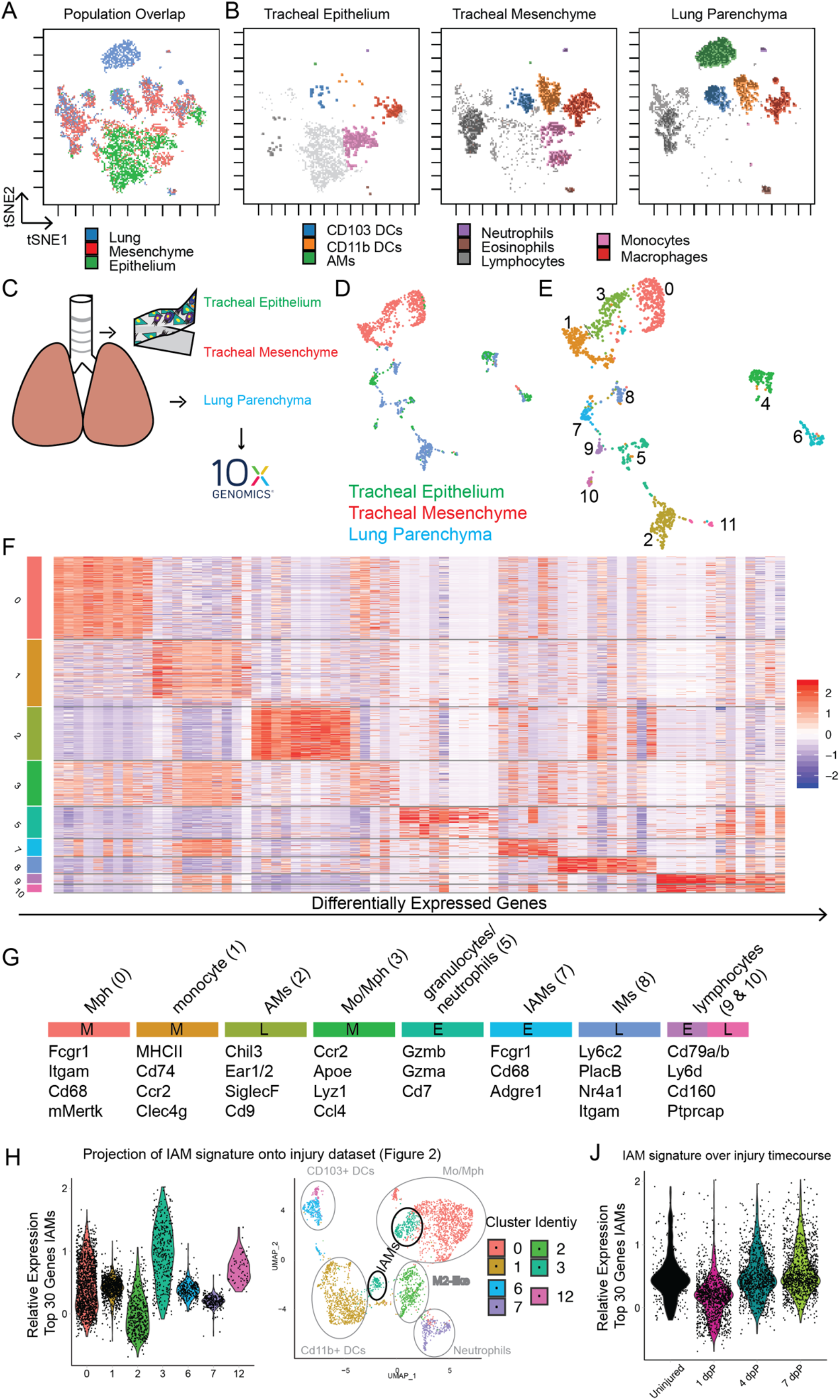
Myeloid cells isolated from the epithelium, mesenchyme and lung parenchyma have distinct transcriptional and translational identities; (A) Unsupervised clustering of 15-marker myeloid panel, FACS analysis of cells from lung, mesenchyme and epithelium (n=5/tissue pooled); (B) Unsupervised clustering of cells isolated from epithelium (left), mesenchyme (middle) and lung (right); (C) Schematic representation of samples isolated for SCSeq, (n=5/tissue animals pooled); captured epithelium=329 cells, mesenchyme= 721 cells, lung= 460 cells; (D) Unsupervised clustering, UMAP, colored for tissue of origin; (E) Unsupervised Louvaine clustering at resolution 0.25 of SCSeq data; (F) Heatmap of differentially expressed genes in myeloid clusters; Clusters 4, 6, and 11 excluded as contaminants for further details see Supplemental Table S3 (G) Cluster assignment based on selected transcripts and annotation of tissue of origin (E – epithelium, M – mesenchyme, L-lung) (H) Projection of IAM gene signature (Cluster 7, Figure 3F) onto single cell injury dataset (Figure 2A-E), (J) Projection of IAM gene signature onto injury time points.

### IAMs, IMs and AMs are molecularly distinct from each other

Because the conventional gating above pointed to unique myeloid populations in the tracheal epithelium and mesenchyme, we performed unbiased SCSeq of CD45^+^ leukocytes isolated from the tracheal epithelium, and CD45^+^;F4/80^+^ cells from the tracheal mesenchyme and lung parenchyma as described above (Figure 3C, S3C). Unsupervised clustering visualized in UMAP space showed that the majority of cells from each tissue formed distinct clusters (Figure 3D). Louvain clustering of these data with a resolution of 0.25 resulted in 12 populations (Figure 3E). Cells in cluster 4 were negative for *Adgre1* and expressed various lymphocyte markers. Cells in cluster 6 and 11 were negative for *Ptprc* (CD45) and *Adgre1*. Cells from Cluster 6 expressed markers associated with fibroblasts (*Col3a1, Clec3b*) and cells in cluster 11 expressed markers of lung epithelium (*Lamp3, Ager*). Therefore clusters 4, 6 and 11 were characterized as non-myeloid cells and excluded from further analysis (Figure S3D, crossed out). We characterized the remaining clusters by identifying the top 10 differentially expressed genes for each (Figure 3F-G, Figure S3E, Supplemental Table S3).

Clusters 0, 1 and 3 were from the tracheal mesenchyme and could be identified as classic macrophages (cluster 0), monocytes (cluster 1) and cells with an intermittent signature (cluster 3) (Figure 3E-G). These three populations are similar to previously identified IMs from the lung, due to their expression of *Mertk, Fcgr1* and *Cd68*, combined with their negativity for *SiglecF* and *Marco* and variable expression of *H2-ab1, H2-aa* and *Retnla* (Gibbings et al., 2017; Ural et al., 2020). A similar signature was also observed for Cluster 8 that originated from the lung, presumably representing the captured lung IMs.

Clusters 5, 7 and 9 were isolated from the tracheal epithelium. Contrary to previous reports (Sertl et al., 1986), these cells do not resemble DCs. Of particular interest was Cluster 7 that contained IAMs. These macrophages are enriched for the expression of *Fcgr1, Cd68* and *Adgre1* (Figure 2G) but also express unique markers such as *Tgfβr1* and *Scimp* (Cluster 7, Supplemental Table S3). Importantly, the transcription profile of these cells was distinct from those reported for IMs (Gibbings et al., 2017), and could not be binned into a classical category. IAMs were also negative for the marker *Cd169*, recently associated with a subpopulation of macrophages localized in close proximity to airway neurons (Ural et al., 2020). Cluster 5 expressed genes associated with neutrophils, and cluster 9 expressed markers of lymphocytes.

Consistent with the flow cytometry data presented above, the cells isolated from the lung parenchyma expressed genes associated with AMs (Cluster 2) and a smaller population of IMs (Cluster 8). Cells in Cluster 10, also isolated from the lung, expressed markers of lymphocytes (Figure 3G, Supplemental Table S3).

For technical reasons, we were unable to separate the epithelium from the underlying mesenchyme after injury prior to SCSeq (Figure 2A-E). Therefore, we derived a transcriptional signature from the top 30 expressed genes in the IAM cluster (Cluster 7 Fig 3E-G, Supplemental Table S4) and analyzed the expression of this signature in the post-injury SCSeq data described above (Figure 2A-E). Cells enriched for this gene signature fell into the monocyte/macrophage cluster (Cluster 3). Consistent with our immunofluorescence data, the IAM gene signature was lowest at 1dpP, when most epithelial cells have died and been sloughed (Figure 3J, Figure S3F). The accumulation of CCR2 lineage traced cells (Figure 2I) concomitant with an increase in the number of cells with an IAM transcriptional signature (Figure 3J and Figure S3F) supports a model in which CCR2^+^ monocytes replace IAMs that are lost as a result of epithelial injury.

### Type 2 cytokines modulate basal cell behaviors *in vitro*

Next, we turned our attention to type 2 cytokines, required for the polarization of M2-like macrophages (Wynn et al., 2013) that increase in the trachea after injury (Figure 2A-C). Our SCSeq of CD45^+^ cells during tracheal injury and repair identified mast cells and group 2 innate lymphoid cells as sources of *Il13* after injury (data not shown). We sought to determine whether these cytokines also directly affect basal cells. We cultured basal cells in the tracheosphere organoid assay (Rock et al., 2009) and supplemented the medium with cytokines associated with lung injury including IL1β, IL2, IL4, IL5 and IL13 (Lechner et al., 2017; Tadokoro et al., 2014) (Figure S4A-B). As previously described (Danahay et al., 2015) the addition of either IL4 or IL13 resulted in an increase in average sphere size (Figure S4A). IL4 and IL13 both signal via a common receptor subunit, IL4Rα. We analyzed a previously published dataset (Plasschaert et al., 2018) and found that *Il4rα* is expressed on some basal cells. Indeed, flow cytometry on isolated basal cells demonstrated that ~11% of basal cells express IL4Rα (Figure 4A). Based on this finding, we tested whether IL4 and IL13 signal directly to basal cells. We infected 2D adherent cultures of IL4Rα ^flox/flox^ basal cells with an adenovirus expressing Cre-GFP fusion protein to delete the floxed *Il4rα* receptor subunit. Three days after infection, Ad5CMVCre-eGFP infected cells, but not AdCMV-GFP controls, had lost IL4Rα expression (Figure S4C). When grown as organoids in the presence or absence of IL4 or IL13 the IL4Rα-deficient cells did not respond to the cytokines, unlike control cells (Figure 4C-D).

**Figure 4:**
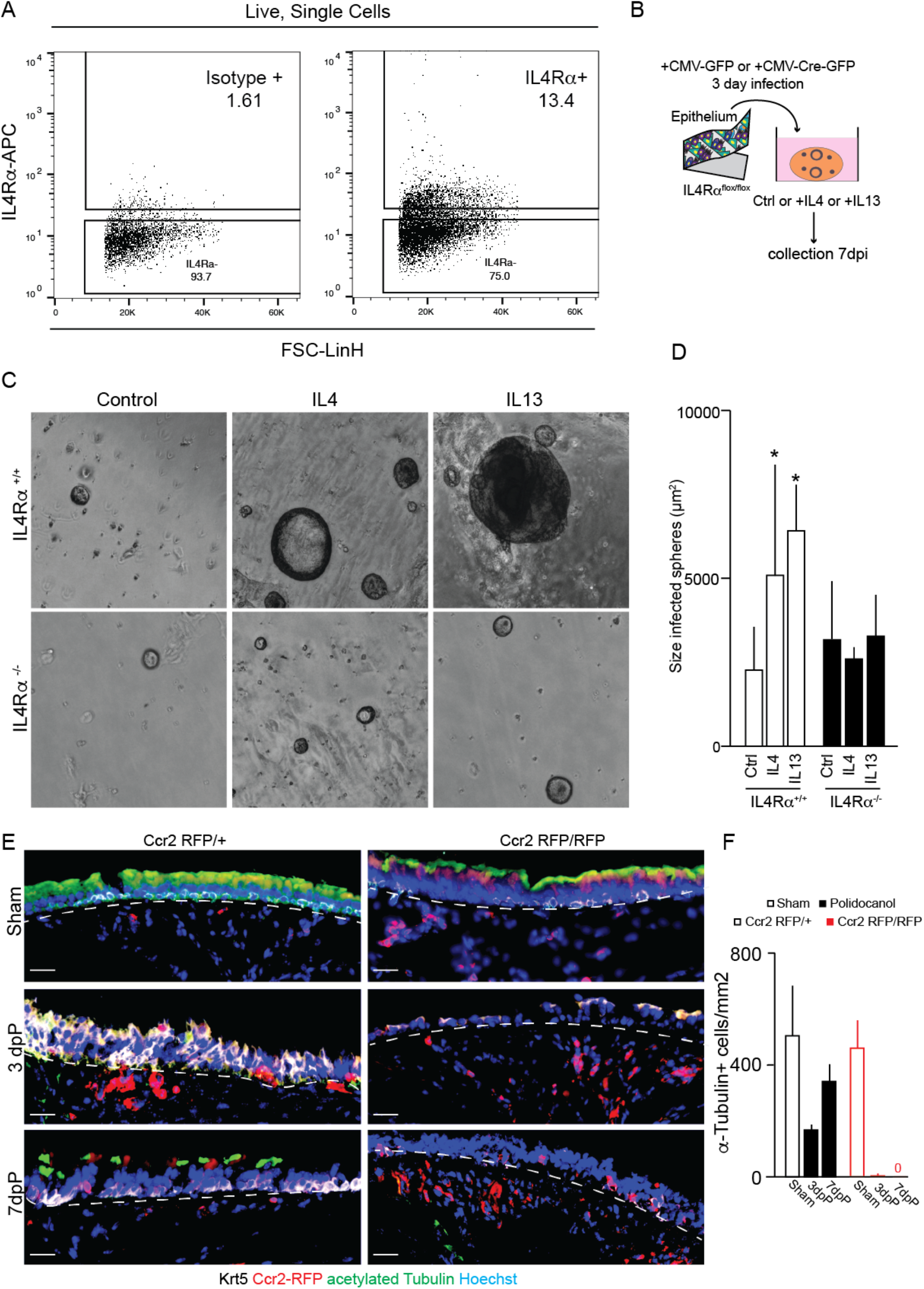
CCR2^+^ myeloid cells promote basal cell mediated epithelial repair. (A) FACS analysis of IL4Rα expression on basal cells (n=3); (B) Schematic representation of basal cell isolation and infection for conditional deletion of IL4Rα and addition of IL4 or IL13 in tracheosphere culture (C) Culture of IL4Rα^+/+^ (n=3) and IL4Rα^-/-^ (n=3) basal cells in the presence of IL4 or IL13; (D) Quantification of sphere size after 10 days of culture; (E) Epithelial repair in CCR2-deficient mice and heterozygous controls 3dpP and 7dpP (heterozygote n(Sham)=2, n(injury)=3 each; homozygote n(Sham) = 4; n(injury)=3 each), scale bars 25 μm and (F) Quantification of acetylated α-Tubulin^+^ ciliated cells as a metric of repair.

### Myeloid cells promote repair of the pseudostratified epithelium

To assess the role of immune cells on tracheal repair *in vivo*, we injured CIAE-NOG animals that lack an adaptive immune system and have a dysfunctional innate immune system including lack of monocytes and macrophages. Three days after polidocanol-induced epithelial injury, basal cells had not undergone extensive self-renewal and temporary stratification, a normal step in the repair process (compare Figure S4D-E to Figure S1D). These data suggest a crucial role of the immune system in fostering epithelial repair. Since there is an accumulation of CCR2 lineage traced cells within the regenerating tracheal epithelium (Figure 2I), we reasoned that CCR2^+^ monocytes and their progeny are required for epithelial repair. To test this hypothesis, we analyzed epithelial repair in mice homozygous for a Ccr2-RFP allele that lack *Ccr2* expression and do not have functioning CCR2^+^ myeloid cells (Saederup et al., 2010) (Figure S4F). Importantly, there were no developmental abnormalities of the tracheal epithelium of Ccr2-RFP homozygotes. Histology and immunofluorescence demonstrated incomplete epithelial repair in *Ccr2*-deficient mice compared to heterozygous littermate controls (Figure 4E-F), characterized by a lack of multiciliated cells.

## Discussion

The mouse trachea is a well characterised model to study airway biology which shares many morphological and histological features with small human airways (Rock et al., 2010). In this study, we used the trachea to interrogate the myeloid cells and their role in homeostasis and epithelial injury and repair. Under steady state conditions we identified macrophages in the tracheal submucosa that are similar to recently described interstitial macrophages (IMs), providing further rationale for our use of the trachea as an airway model. We unexpectedly identified a population of macrophages within the tracheal epithelium that are molecularly distinct from IMs and AMs. These intraepithelial macrophages (IAMs) are also distinct from DCs which have been described within the epithelium (Sertl et al., 1986). Similarly, Langerhans cells, previously considered a specialized dendritic cell of the epidermis, have recently been reclassified as a tissue resident macrophage population (Doebel et al., 2017). Under baseline conditions, both IAMs and Langerhans cells are quiescent, defined by negativity for cell cycle markers (*Ccne2, Aurka, Mki67*) in SCSeq, absence of Ki67 by immunostaining (data not shown), and directly contact surrounding epithelial cells (Clayton et al., 2017; Doebel et al., 2017).

Our flow cytometry and SCSeq of CD45^+^;F4/80^+^ cells revealed several dynamic properties of trachea-associated myeloid populations in response to epithelial injury. For example, at 1 and 3dpP we detected an increase in neutrophils, monocytes and macrophages. Neutrophils are thought to be key in modulating the cytokine environment to foster recruitment and differentiation of other immune players (Blázquez-Prieto et al., 2018; Kaplanski et al., 2003; Lechner et al., 2017; Mindt et al., 2018; Ricardo-Gonzalez et al., 2018). In the monocyte and macrophage compartment, we observed a shift toward M2-like polarization of macrophages. We confirmed the accumulation of Arginase-expressing M2-like macrophages after epithelial injury using transgenic reporter mice.

By combining the Arginase reporter allele with a Ccr2-based lineage tracing strategy, we provide evidence that some of the M2-like macrophages come from a Ccr2-expressing precursor. Similar to previous reports, we also observed Cx3cr1-expressing monocytes in homeostatic tracheas and in response to injury (Joshi et al., 2020; Ural et al., 2020). Whether or not these cells also give rise to M2-like macrophages and IAMs remains unknown. The accumulation of CCR2-derived cells concomitant with the reemergence of the IAM transcriptional signature 4 and 7 dpP suggests that CCR2^+^ monocytes are a precursor for IAMs in response to injury.

Monocyte-derived, alternatively activated, M2-like macrophages are crurcial for development, regeneration and remodeling in disease (Byrne et al., 2015; Lechner et al., 2017). Our previous work demonstrated the importance of M2-like macrophages for the compensatory regrowth of alveoli in response to pneumonectomy in mice (Lechner et al., 2017). We used parallel strategies to determine whether leukocytes also promote the repair of the tracheal epithelium. First, we performed polidocanol-injury of the airway epithelium in immunodeficient mice and demonstrated a deficiency in basal cell proliferation and differentiation 3 days post-injury. Second, we analyzed tracheal repair in Ccr2-deficient mice to more specifically determine whether monocytes and their progeny are required for epithelial repair. Similar to immunodeficient mice, there was inefficient repair of the tracheal epithelium in Ccr2-deficient mice, characterized by failure to generate a pseudostratified epithelium with multiciliated cells beginning 3 days postinjury.

Our SCSeq data of CD45^+^;F4/80^+^ cells revealed that the type 2 cytokines, IL4 and IL13, are expressed by group 2 innate lymphoid cells and mast cells in the trachea (data not shown). In addition to numerous other functions, these factors are critical for the polarization of M2-like macrophages. Future experiments will identify the relative contributions of type 2 cytokines by mast cells, ILC2s and other non-myeloid lineages. Because we detected IL4 and IL13 in the injured trachea, we sought to confirm previous reports that these cytokines directly modulate epithelial basal cell behaviors (Danahay et al., 2015; Kuperman et al., 2002). The treatment of wild type basal cells (but not *Il4rα*-deficient basal cells) with IL4 or IL13 resulted in an increased size of tracheospheres. The cellular basis for this increase remains to be determined, but highlights the pleotropic effects of type 2 cytokines during repair of the tracheal epithelium.

This work and recent work by others (Chakarov et al., 2019; Gibbings et al., 2017; Lechner et al., 2017; Misharin et al., 2013; Ural et al., 2020) illustrate the striking heterogeneity within the myeloid compartment of the respiratory system. With modern, single cell technologies we can slowly appreciate the high complexity of tissue resident monocytes and macrophages which will be key for the development of future, targeted technologies and therapies (Klichinsky et al., 2020).

## Contact for Reagents

This study did not generate new, unique reagents. Datasets are available in repositories. Further information and requests for resources and reagents should be directed to, and will be fulfilled by, the Lead Contact, Jason R. Rock.

## Author Contributions

Conceptualization: AEE, SAM, JRR; Methodology and Investigation AEE, ABY, RMFP, ACB, HH, NM, and HR; Data Analysis and Interpretation: AEE, CVM, ACB and JRR; Writing, AEE and JRR, Writing – Review and Editing; AEE, ABY, ACB, SAM and JRR; Supervision, Project Administration, and Funding Acquisition, SAM and JRR.

## Acknowledgments

We would like to thank the entire Rock lab for helpful discussions. We thank Drs. Darrell Kotton, Brigid Hogan and Jay Mizgerd for helpful comments on the manuscript. We thank Brian Tilton of the Boston University School of Medicine Flow Cytometry core Facility, Dr. Yuriy Alekseyev and Ashley LeClerc of the Boston University School of Medicine Sequencing core and Dr Michael T. Kirber Boston University School of Medicine Cellular imaging core for excellent technical assistance.

