## Supplemental Materials for "Airway-associated macrophages in homeostasis and repair"

Experimental Procedures

Supplemental Figure S1
Supplemental Figure Legend S1
Supplemental Figure S2
Supplemental Figure Legend S2
Supplemental Figure S3
Supplemental Figure Legend S3
Supplemental Figure S3
Supplemental Figure Legend S3

Supplemental Table S1 – 15-color myeloid panel
Supplemental Table S2 – SCSeq Injury dataset (Accessible after Publication)
Supplemental Table S3 – SCSeq Tissue dataset (Accessible after Publication)
Supplemental Table S4 – Gene Sets for Enrichment Analyses

**Experimental Procedures (Star Methods)**

**KEY RESOURCES TABLE**

| REAGENT or RESOURCE | SOURCE | IDENTIFIER |
|---|---|---|
| Antibodies | | |
| CD45, BUV395 (clone 30-F110) | BD Bioscience | Cat#564279 |
| CD19, BUV737 (clone ID3) | Becton Dickinson | Cat#612781 |
| CD64, BV421 (clone X54-5/7.1) | Biolegend | Cat#139309 |
| CD24, BV510 (clone M1/69) | Biolegend | Cat#101831 |
| CD80, BV605 (clone 16-10a1) | Biolegend | Cat#104729 |
| CD86, BV650 (clone 24F) | BD Bioscience | Cat#743214 |
| MHC II (I-A/I-E), FITC (clone M5/114.15.2) | Biolegend | Cat#107606 |
| CD40, PE (clone 3/23) | BD Bioscience | Cat#561846 |
| Siglec F, PE-CF594 (clone E50-2440) | BD Bioscience | Cat#562757 |
| Ly6C, PerCp5.5 (clone HK1.4) | Biolegend | Cat#128012 |
| F4/80, PE-Cy7 (clone BM8) | Biolegend | Cat#123114 |
| CD103, APC (clone 2E7) | Biolegend | Cat#121432 |
| Ly6G, AF700 (clone 1A8) | BD | Cat#127622 |
| CD11b, APCFire (clone M1/70) | Biolegend | Cat#101262 |
| CD45, APC (clone 30-F11) | Biolegend | Cat#103112 |
| CD45, FITC (clone 30-F11) | Biolegend | Cat#103108 |
| CD45, BV421 (clone 30-F11) | Biolegend | Cat#103134 |
| CD45.2, BUV737 (clone 104 ) | BD Bioscience | Cat#612778 |
| F4/80, APC (clone BM8) | Biolegend | Cat#123116 |
| F4/80, BV421 (clone BM8) | Biolegend | Cat#123132 |
| CD11b, FITC (clone MI/70) | Biolegend | Cat#101206 |
| CD11c, PeCy7 (clone N418) | Biolegend | Cat#117317 |
| SiglecF, Alexa647 (clone E50-2440) | BD Bioscience | Cat#562680 |
| CD64, PE (clone X54-5/7.1) | Biolegend | Cat#139303 |
| Ly6G, PECy7 (clone 1A8) | Biolegend | Cat#127617 |
| IL4Ra, APC (clone I015F8) | Biolegend | Cat#144807 |
| Krt5 (rabbit) | Biolegend | Prb-160P |
| Acetylated tubulin (mouse) | Sigma | T7451 |
| F4/80 (rat) | Biorad | MCA497GA |
| RFP (goat) | MyBioSource | MBS448122 |
| CD45 (goat) | R&D | AF114 |
| Bacterial and Virus Strains | | |
| Ad-CMV-GFP | University of Iowa Viral Vector Core | Ad5CMVeGFP |
| Ad-CMV-Cre-GFP | University of Iowa Viral Vector Core | Ad5CMVCre-eGFP |
| Chemicals, Peptides, and Recombinant Proteins | | |
| Tamoxifen | Sigma Aldrich | Cat#T5648 |
| Polidocanol | Sigma Aldrich | Cat#88315-100G |
| “Isothesia” Isoflurane | Henry Schein | Cat#029405 |
| Normal Donkey Serum | Jackson ImmunoResearch | Cat# 017-000-121 |
| Bovine Serum Albumin | Fisher Chemical | Cat#BP1600-100 |

| | | |
|---|---|---|
| Triton-X | Fisher BioReagents | Cat#BP151-100 |
| EdU (5-ethynyl-20-deoxyuridine) | Invitrogen | Cat#C10340 |
| Collagenase A (1.5 mg/mL) | Roche | Cat#10103586001 |
| DNase I (0.4 mg/mL) | Roche | Cat#10104159001 |
| Dispase | Corning | Cat#354235 |
| RPMI 1640 Media | Gibco | Cat#11875093 |
| Hoechst 33342p | Invitrogen | Cat#H1399 |
| Calcein Blue, AM | Invitrogen | Cat#C1429 |
| D-Sucrose | Fisher Bioreagents | Cat#BP220-1 |
| OCT Compound | Fisher Healthcare | Cat#4585 |
| Paraffin | Fisher Chemical | Cat#T565 |
| Xylenes | Fisher Chemical | Cat#X3S-4 |
| Paraformaldehyde | Fisher Chemical | Cat#O4042-500 |
| Antigen Unmasking Solution, Citric Acid Based | Vector Labs | Cat#H-3300 |
| DMEM/F-12 | Gibco | Cat#11330-057 |
| HEPES | Fisher Bioreagents | Cat#BP299-1 |
| Sodium Bicarbonate (7.5%) | Sigma-Aldrich | Cat#S8761 |
| L-Glutamine | Gibco | Cat#25030164 |
| Pen-Strep | Gibco | Cat# 15140-122 |
| Amphotericin B | Gibco | Cat#152-90-018 |
| Insulin-Transferrin-Selenium | Gibco | Cat#41400045 |
| Cholera Toxin | Sigma-Aldrich | Cat#C8052 |
| Recombinant Mouse Epidermal Growth Factor | R&D Systems | Cat#2028-EG-200 |
| Bovine Pituitary Extract | Gibco | Cat#13028-014 |
| Trypsin (0.25%) | Gibco | Cat#25200114 |
| Retinoic Acid | Sigma-Aldrich | Cat#R2625 |
| GFR 3D Matrigel | Corning | Cat#356231 |
| Recombinant Mouse Il1b Protein | R&D Systems | Cat#401-ML-010 |
| Recombinant Mouse IL2 Protein | R&D Systems | Cat#402-ML-020 |
| Recombinant Mouse IL4 Protein | R&D Systems | Cat#404-ML-010 |
| Recombinant Mouse IL5 Protein | R&D Systems | Cat#405-ML-005 |
| Recombinant Mouse IL13 Protein | R&D Systems | Cat#413-ML-005 |
| Experimental Models: Organisms/Strains | | |
| Tg(Csf1r-EGFP)1Hume/J mouse | The Jackson Laboratory, made by Sasmono et al 2003 | Stock#018549 |
| Ccr2$^{tm2.1Ifc}$/J mouse, heterozygous and homozygous | The Jackson Laboratory, Made by Saederup et al 2010 | Stock#017586 |
| B6.129S4-Arg1$^{tm1Lky}$/J mouse | The Jackson Laboratory, made by Reese et al 2007 | Stock#015857 |
| Il4ra$^{tm1Sz}$/J mouse | Received from NIH, made by Noben-Trauth et al 1997 | Stock#003514 |
| Software and Algorithms | | |
| Fiji/ImageJ | Fiji | https://fiji.sc/ |

| | | |
|---|---|---|
| NIS-Elements | Nikon | https://www.microscope.healthcare.nikon.com/products/software/nis-elements |
| Leica Application Suite X | Leica | https://www.leica-microsystems.com/products/microscope-software/p/leica-las-x-ls/ |
| FlowJo | BD Biosciences | https://www.flowjo.com/ |
| Cytobank | Cytobank | https://www.cytobank.org/ |
| Cell Ranger | 10X Genomics | v.2.0.1 |
| Seurat | Satija Lab | v3 |
| OMIQ | OMIQ | https://www.omiq.ai |
| Other | | |
| Cryostat | Leica | CM1950 |
| Inverted fluorescent microscope | Nikon | Eclipse NiE |
| Confocal | Leica | SP6 |
| Flow Cytometer (Sorting) | Beckman Coulter | MoFlo Astrios |
| Flow Cytometer (Analysis) | BD Biosciences | Aria II |
| Stratedigm | Stratedigm | S1000EXI |

**Animal Husbandry**

Experiments were conducted as sex unbiased, with a minimum of three animals per experimental group. All mice were bred and maintained in a specific-pathogen-free barrier facility with free access to food and water. Both male and female mice were used between 8-12 weeks of age for all experiments. All studies were approved and performed according to the guidelines issued by Boston University's (BU) Institutional Animal Care and Use Committees (IACUC) under license numbers PROTO201900002. Animals were commercially acquired via Jackson Laboratories. Tg(Csf1r-EGFP)1Hume/J (also referred to as *Csf1r-GFP*) mice (Sasmono et al., 2003), *Ccr2*$^{tm2.1Ifc}$ (also referred to as *Ccr2*$^{RFP/+}$, (Saederup et al., 2010)) mice were used for macrophage and monocyte studies and *Ccr2*$^{RFP/RFP}$ (pure-bred C57BL/6 strain) mice were used to assess the contribution of Ccr2$^{+}$ cells after Polidocanol mediated injury. *Arg1*$^{tm1Lky/J}$ ((Reese et al., 2007) also referred to as YARG) mice were used to assess M2-like polarization of macrophages and *Il4ra*$^{tm1Sz}$ mice were used to isolate basal cells for culture assays.

**Tamoxifen Administration**

Tamoxifen (Sigma-Aldrich, catalog #T5648) was dissolved in corn oil to a final concentration of 20mg/mL and administered to mice via intraperitoneal injection at 0.25 mg of Tamoxifen per kg of mouse weight. A single full dose of TMX was given to each animal. Further analysis and additional experiments were performed earliest 4-days after last Tamoxifen dosage (early time-points were used for replenishment studies) and for progenitor analysis additional experiments were performed after full Tamoxifen washout after 14 days (YARG coexpression experiments.

**Polidocanol Induced Injury**

Polidocanol-induced injury was performed as previously described (Borthwick 2001). Mice were anesthetized in an Isoflurane chamber and delivered one dose of 15 µL freshly prepared, 2% Polidocanol or PBS sham control by oropharyngeal aspiration to induce injury. Tracheas were harvested 1-, 3- or 7-days following injury for scRNA-Seq, FACS Analysis or immunohistochemistry.

**Tissue Preparation**

Mice were euthanized, chest cavity was opened and tracheas exposed. For scRNA-seq experiments the pulmonary vasculature was perfused via the right heart ventricle. Uninjured tracheas were peeled as

previously described (Rock et al., 2009), in short, tracheas were dissected, cleaned, cut in smaller pieces and predigested in Dispase (15 U/mL, Sigma-Aldrich) for maximally 25 minutes and the epithelium was thereafter peeled from the mesenchyme. The epithelium was further digested in trypsin (0.25%) for 15 minutes.
Whole tracheas, leftover mesenchyme and lung were digested with 1.5 mg/mL Collagenase A (Roche), 0.4 mg/mL DNase I (Roche), and 2 U/mL Dispase (Sigma–Aldrich) in RPMI base medium at 37 °C for 30-45 minutes. Reaction was stopped using FBS to a concentration of 10%.
For tissue collection for immunohistochemistry, whole throats were dissected and fixed for maximally 4 hours in 4% PFA. PFA was washed out of tissue using PBS and the trachea was dissected out of the throat. Tracheas were either embedded for Paraffin histology, by moving through ascending Ethanol (50%-100%), Xylene and Paraffin in a vacuum oven and sectioned at 5-7 µm on a microtome (Leica) or were prepared for frozen sectioning, by cryopreservation in 30% Sucrose for 48 hours and embedding in OCT medium and freezing on dry ice. Frozen organs were sectioned on Cryostat (Leica) at 10-12 µm thickness within one month and sectioned tissue was stored at −20°C.

**Tracheosphere Culture**
Basal Cells were Cultured as previously described (Rock et al., 2009). In brief, basal cells were suspended in mouse tracheal epithelial cells (MTEC)/plus medium mixed at a 1:1 ratio with growth factor-reduced Matrigel (BD Biosciences), and seeded at 500 cells/90 µL droplet. Medium was changed every other day. Analysis was done on two independent experiments using three independent primary cultures. For IL4R□ deletion, primary cells from IL4R□$^{flox/flox}$ animals were isolated and infected as pure, adherent basal cell cultures using MOI 3, Ad-CMV-GFP (Control) or Ad-CMV-Cre-GFP (Cre) virus (Viral Vectors were provided by the University of Iowa Viral Vector Core (http://www.medicine.uiowa.edu/vectorcore). 3-days after infection cells were dissociated from plastic and analysed for deletion by FACS and plated as previously described for differentiation as tracheosphere.

**Immunohistochemistry**
Paraffin slides were rehydrated by consecutive, descending processing through Xylene and Ethanol (100%-50%). Paraffin slides underwent citrate-based retrieval (Vector Unmasking), for frozen sections only select panels were citrate retrieved.
Tissue sections were blocked and permeabilized for 30 – 60 minutes in 10% NDS, 4% BSA and 0.5% TritonX. Primary antibodies (Krt5 (rb) Covance (Prb-160P) 1:1000; F4/80 (rt) Biorad (MCA497GA) 1:250; RFP (gt) Abcam (ab25877) 1:500; acet.Tub (ms) Sigma (T7451) 1:2000; CD45 (gt) R&D (AF114) 1:250) were diluted in primary block solution (2.5% NDS, 1% BSA, 0.125% TritonX) and incubated overnight at 4°C in a humidified chamber. All fluorophore-conjugated secondary antibodies were diluted in secondary block solution (5% NDS, 2% BSA, 0.25% TX) at 1:500 containing Höechst nuclear stain dye. EdU staining was performed according to manufacturer recommendations (Invitrogen). Images of sections were captured on a Nikon Eclipse NiE or a Leica SP6.

**Flow Cytometry**
Whole tissue digests to obtain single cell suspension (as described in tissue preparation) were kept on ice and stained in flow cytometry/FACS buffer (1% BSA in PBS without $Ca^{2+}$ or $Mg^{2+}$). For live cell staining and sorting, a cell viability dye (DAPI or Calcein Blue) was added before sample acquisition. Flow cytometry analysis was performed on a Stratadigm (S1000EXI) or FACSAria II (BD Biosciences) and cell sorting was performed on Moflo Astrios (Beckman Coulter). Compensation was performed with single-stained UltraComp compensation beads (Thermo Fisher) or single-stained immune cells prepared from spleen or lung. To determine gating controls for reporter gene expression, mice lacking the reporter or protein were used. Initial data cleanup and expert-driven analysis were performed using FlowJo (BD Biosciences) and Cytobank cloud-based analysis software (Beckman Coulter). For the 15-color myeloid cell immunophenotyping, intravascular immune cells were labelled by tail vein injection (Anderson et al., 2014) of 2 µg of CD45-BUV737 (BD Biosciences) diluted in 100 µL of sterile saline and loaded into a 28.5 gauge insulin syringe. For tail vein injection animals were lightly anesthesized, injected and kept under controlled, anesthesized conditions, before right ventricle transcardial perfusion 3 minutes post injection. Automated analysis of data generated from representative samples was performed in Omiq.ai cloud-based analysis platform. CD45+ live single cells phenotyped across 15 markers were clustered with Phenograph unsupervised clustering algorithm (Levine et al., 2015) and projected into opt-SNE space (Belkina et al.,

2019). Median fluorescence intensities of each marker across Phenograph-found populations were plotted on a hierarchically cluster heatmap to allow expert annotation of specific populations.

**Single Cell Sequencing**

Tissue was prepared as described above for fluorescent activated cell sorting (FACS). For mesenchymal and lung samples viable $CD45^+$;$F4/80^+$ cells were sorted, for peeled tracheal epithelium and tracheas in Polidocanol injury experiments viable bulk $CD45^+$ cells were sorted and brought to appropriate concentration, according to 10X recommendation for capture of up to 2000 cells/lane. Single cells were captured for sequencing library preparation using a 10X Chromium (10X Genomics, Pleasanton, CA) instrument at the BUMC Single Cell Core. Single-cell RNA-seq libraries were prepared according to the Single Cell 3' v2 Reagent Kits User Guide (10X Genomics). Sequence libraries were constructed using the Chromium Single-Cell 3' Library Kit (10X Genomics). Sequencing libraries were loaded on a NextSeq500 (Illumina) to obtain a sequencing depth of~200K reads per cell. 329 cells from the tracheal epithelium, 721 cells from the tracheal mesenchyme and 460 cells from whole lung parenchyma were captured and sequenced. For the epithelium 143,824 mean reads per cell were obtained corresponding to 2133 UMI counts and 908 genes expressed per cell. For the mesenchyme 190,186 reads per cell were obtained corresponding to 5383 UMI counts and 1891 genes expressed per cell. For the lung parenchyma, 320,071 mean reads per cell were obtained corresponding to 3969 UMI counts per cell and 1636 genes expressed per cell.

**Single Cell Analysis**

Single cell reads were mapped to the mouse genome reference (ENSEMBL, GRCm38) and pre-processed with Cell Ranger v.2.0.1 to obtain the matrix of UMI counts per gene per cell. We used Seurat v.3 to normalize, scale and regress out unwanted sources of variation (like cell degradation), and subsequently identified highly variable genes for linear dimensionality reduction with principal component analysis (PCA). The principal components were then used for clustering (using the Louvain algorithm). Further non-linear dimensionality reduction was done with Uniform Manifold Approximation and Projection (UMAP) for visualization purposes. Differential gene expression was tested using MAST (Finak et al., 2015). Clusters identified with the Louvain method were annotated based on their differentially expressed genes (DEGs).

**Quantification and Statistical Analysis**

Stained sections were analyzed with a Nikon Eclipse NiE or a Leica SP6. Images were processed with Fiji/ImageJ. For quantification of cells, at least 3 randomly selected, non-consecutive regions per trachea were imaged and counted per animal. Images were processed with ImageJ/FIJI (version 2.0.0, NIH). Statistical significance was determined by two-tailed Student's t test on mean values, and percentages and proportions were arcsine square root transformed before statistical testing. Significance was determined as $^*p < 0.05$, $^{**}p < 0.01$, and $^{***}p < 0.001$; Deviance from the mean is displayed as standard deviation.

Figure S1

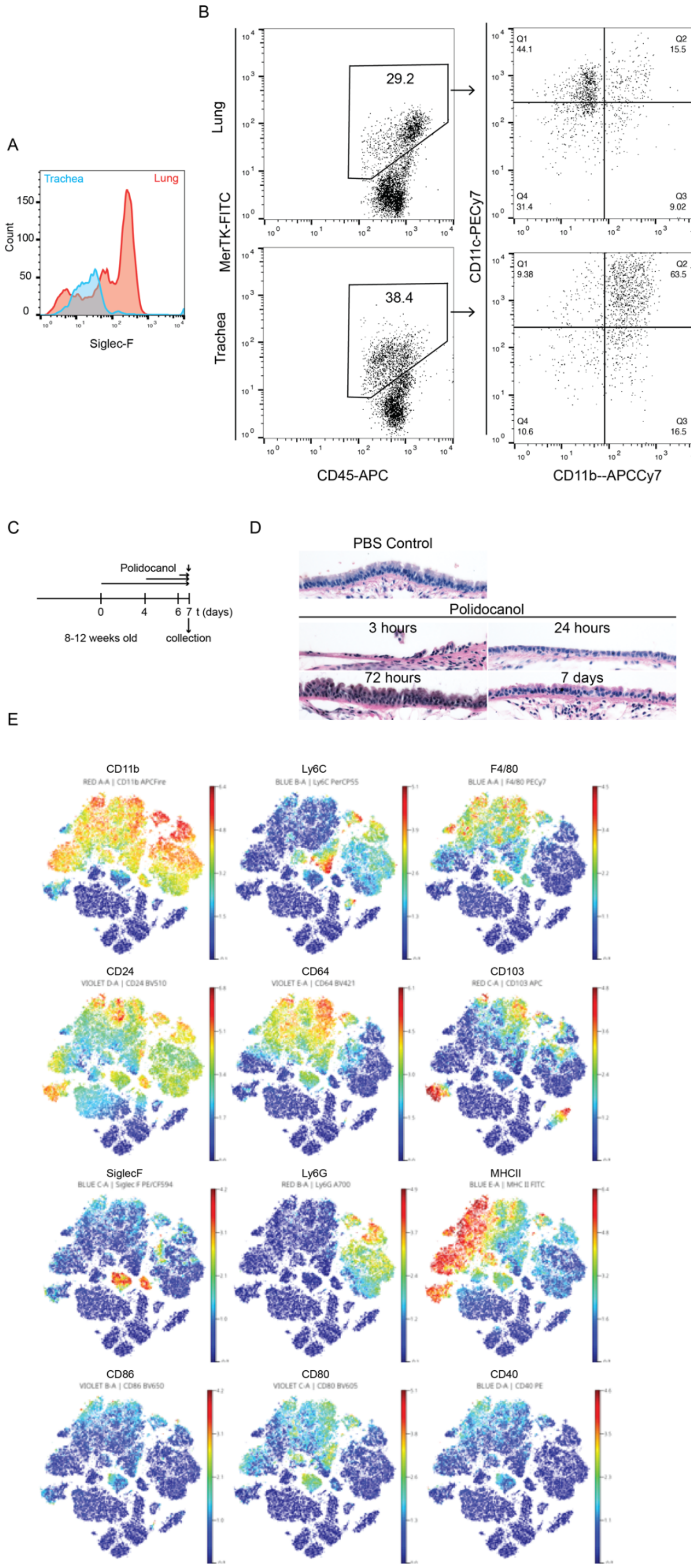
A
Trachea
Lung
Count
Siglec-F
B
Lung
Trachea
MerTK-FITC
CD45-APC
29.2
38.4
CD11c-PECy7
CD11b--APCCy7
Q1 44.1
Q2 15.5
Q4 31.4
Q3 9.02
Q1 9.38
Q2 63.5
Q4 10.6
Q3 16.5
C
Polidocanol
0
4
6
7
t (days)
8-12 weeks old
collection
D
PBS Control
Polidocanol
3 hours
24 hours
72 hours
7 days
E
CD11b
Ly6C
F4/80
CD24
CD64
CD103
SiglecF
Ly6G
MHCII
CD86
CD80
CD40

**Figure S1: Temporal changes of the tracheal innate myeloid compartment;** (A) FACS Analysis of whole trachea digest versus lung at baseline; gated on CD45+F4/80+ cells, SiglecF expression (n=3) (B) FACS Analysis of whole trachea digest versus lung at baseline; Macrophage gating on $CD45^{+}$;$MerTK^{+}$ cells, with expression analysis of CD11b and CD11c; (n=3) (C) Schematic representation of Polidocanol mediated injury timeline (D) Histology of Polidocanol mediated injury 3hrs, 1dpP, 3dpP and 7dpP (E) Sham, 1dpP, 3dpP and 7 dpP myeloid cell data concatenated and graphed as opt-SNE dimensionality reduction plots with individual marker median expression projected as a heatmap (n=1/time point, pooled 2 tracheas per sample).

Figure S2

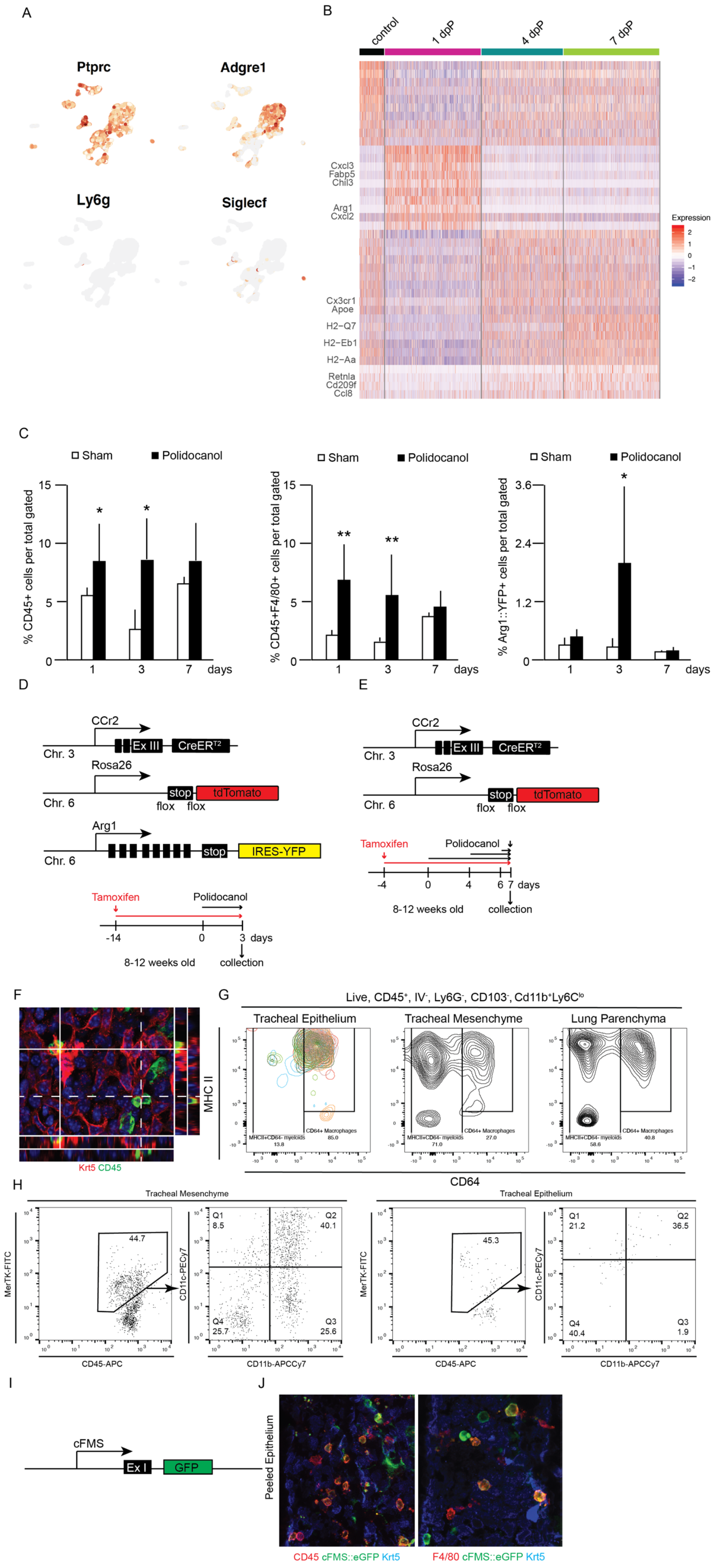
A
Ptprc
Adgre1
Ly6g
Siglecf
B
control
1 dpP
4 dpP
7 dpP
Cxcl3
Fabp5
Chil3
Arg1
Cxcl2
Cx3cr1
Apoe
H2−Q7
H2−Eb1
H2−Aa
Retnla
Cd209f
Ccl8
Expression
C
Sham
Polidocanol
% CD45+ cells per total gated
% CD45+F4/80+ cells per total gated
% Arg1::YFP+ cells per total gated
days
D
CCr2
Ex III
CreER
Chr. 3
Rosa26
stop
tdTomato
flox
Chr. 6
Arg1
IRES-YFP
Tamoxifen
Polidocanol
8-12 weeks old
collection
E
F
Krt5 CD45
G
Live, CD45+, IV-, Ly6G-, CD103-, Cd11b+Ly6Clo
Tracheal Epithelium
Tracheal Mesenchyme
Lung Parenchyma
MHC II
CD64
H
Tracheal Mesenchyme
Tracheal Epithelium
MerTK-FITC
CD11c-PECy7
CD45-APC
CD11b-APCCy7
I
cFMS
Ex I
GFP
J
Peeled Epithelium
CD45 cFMS::eGFP Krt5
F4/80 cFMS::eGFP Krt5

**Figure S2: Unbiased analysis of tracheal myeloid populations after injury;** (A) SCSeq after injury, select gene expression (Ptprc/CD45; Adgre1; Ly6G, SiglecF); (B) Heatmap top differentially expressed genes after injury (n=1, pooled 3 tracheas); (C) Quantifications of FACS analysis of injury time line %CD45+ cells, %CD45+F4/80+ cells and % total YARG+ cells; (1dpP n=3; 3dpP n(Sham)=4, n(Poli = 3); 7dpP: n(Sham)=4, n(Poli)=6); (D) Graphical representation of Ccr2-CreER$^{T2}$;ROSA-tdTom, YARG triple transgenics and schematic representation of Tamoxifen and Polidocanol timeline; (E) Graphical representation of Ccr2-CreERT2;ROSA-tdTom YARG double transgenics and schematic representation of Tamoxifen and Polidocanol timeline; (F) peeled epithelium Krt5 red, CD45 green, nuclear stain Höechst; (G) FACS analysis of Live, CD45$^{+}$, extravascular, Ly6G$^{-}$, CD103$^{-}$, CD11b$^{+}$, Ly6C$^{lo}$ cells for MHCII, CD64 expression; (n=3); (H) FACS Analysis of mesenchyme and peeled epithelium; Macrophage gating on CD45$^{+}$; MertK$^{+}$ cells, with expression analysis of CD11b and CD11c (n=3); (I) Graphical representation of cFMS::eGFP transgenic animals; (J) peeled epithelium of cFMS::eGFP animals; Krt5 blue, cFMS::eGFP green, CD45 red (left) and F4/80 red (right) scale bar 25 μm;

Figure S3

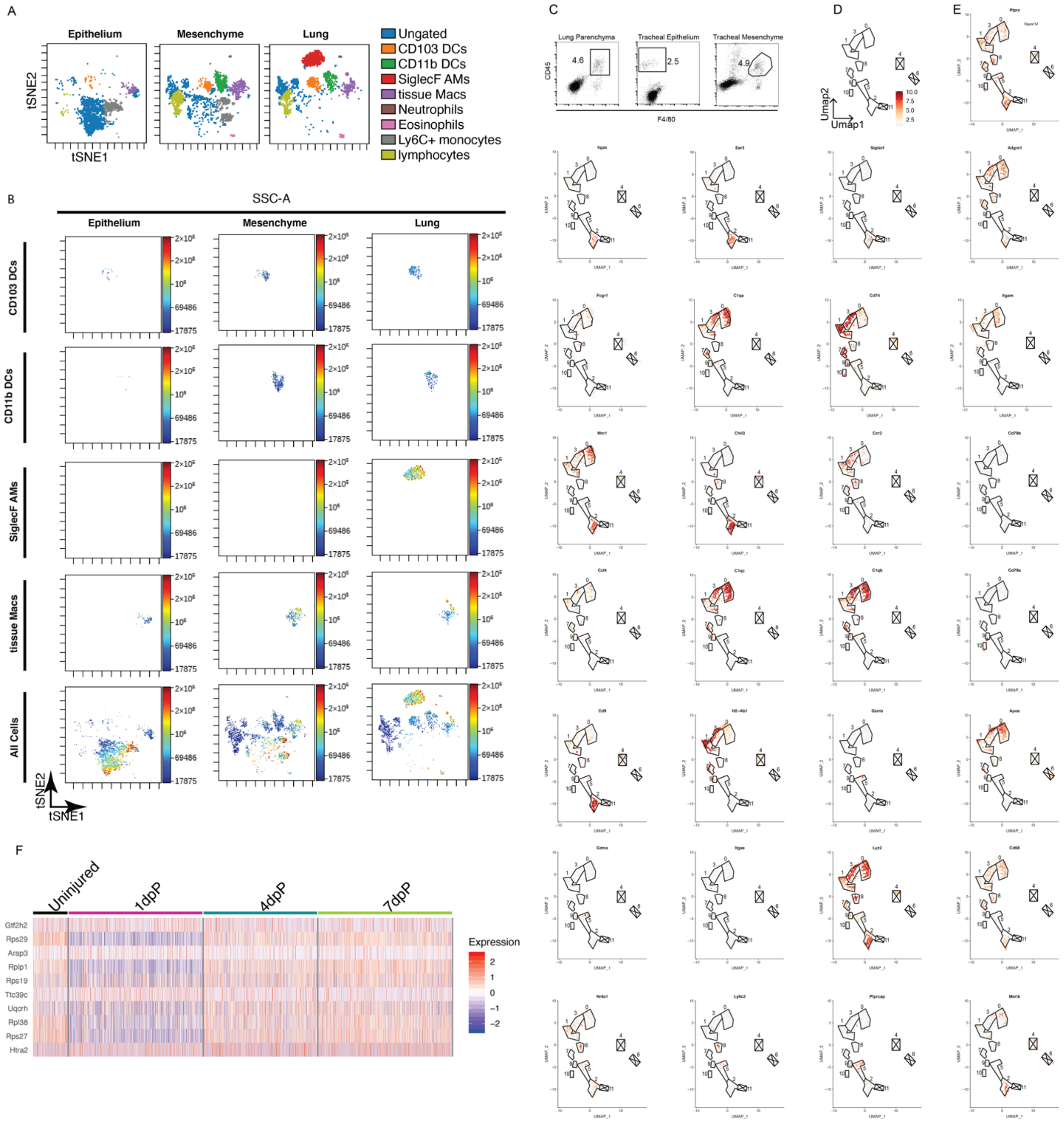

A
Epithelium
Mesenchyme
Lung
tSNE2
tSNE1
Ungated
CD103 DCs
CD11b DCs
SiglecF AMs
tissue Macs
Neutrophils
Eosinophils
Ly6C+ monocytes
lymphocytes
B
SSC-A
Epithelium
Mesenchyme
Lung
CD103 DCs
CD11b DCs
SiglecF AMs
tissue Macs
All Cells
2×10^6
10^6
69486
17875
tSNE2
tSNE1
C
Lung Parenchyma
Tracheal Epithelium
Tracheal Mesenchyme
CD45
F4/80
4.6
2.5
4.9
D
Umap2
Umap1
10.0
7.5
5.0
2.5
E
F
Uninjured
1dpP
4dpP
7dpP
Gtf2h2
Rps29
Arap3
Rplp1
Rps19
Ttc39c
Uqcrh
Rpl38
Rps27
Htra2
Expression
2
1
0
−1
−2

**Figure S3: Myeloid cells isolated from the epithelium or the mesenchyme or the lung parenchyma have different transcriptional and translational identities;** (A) Unsupervised clustering of 15-color myeloid panel lung, mesenchyme and epithelium (n=5 pooled); **(**B) Individual marker expression per tissue of origin of 15-marker FACS panel and expert driven gating; (C) FACS sorting of myeloid cells for single cell sequencing; (D) Schematic representation of clusters; Unsupervised Louvain clustering resolution 0.25, clusters 4, 6 and 11 excluded (n=5 pooled/sample); (E) Unsupervised Louvaine clustering resolution 0.25; of single cell sequencing, indentifying genes for expert driven cluster naming, from top to bottom, left to right: Ptprc, Itgax, Ear2, SiglecF, Adgre1, Fcgr1, C1qa, Cd74, Itgam, Mrc1, Chil3, Ccr2, Cd79b, Ccl4, C1qc, C1qb, Cd79a, Cd9, H2-Ab1, Gzmb, Apoe, Gzma, Itgae, Lyz2, Cd68, Nr4a1, Ly6c2, Ptprcap, Mertk; (F) Heatmap of genes in Cluster 3 of injury dataset, analysis changes in IAM dynamics upon injury (n=3 pooled/sample).

Figure S4

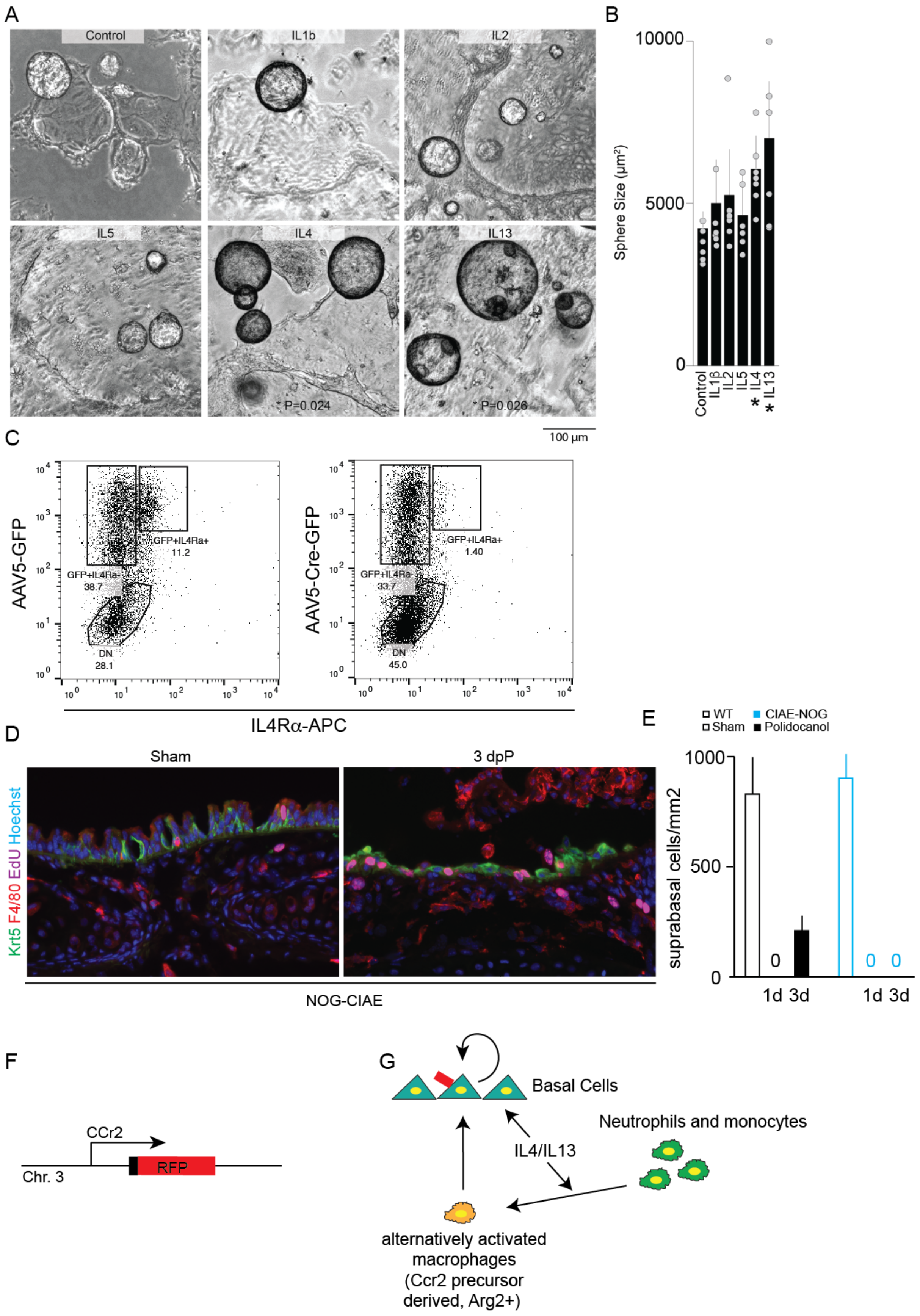

A
Control
IL1b
IL2
IL5
IL4
IL13
P=0.024
P=0.026
100 µm
B
Sphere Size (µm²)
10000
5000
0
Control
IL1β
IL2
IL5
IL4
IL13
C
AAV5-GFP
AAV5-Cre-GFP
GFP+IL4Ra+
11.2
GFP+IL4Ra-
38.7
DN
28.1
GFP+IL4Ra+
1.40
GFP+IL4Ra-
33.7
DN
45.0
IL4Rα-APC
D
Sham
3 dpP
Krt5 F4/80 EdU Hoechst
NOG-CIAE
E
WT
Sham
CIAE-NOG
Polidocanol
suprabasal cells/mm2
1000
500
0
0
0
0
1d 3d
1d 3d
F
CCr2
RFP
Chr. 3
G
Basal Cells
Neutrophils and monocytes
IL4/IL13
alternatively activated macrophages (Ccr2 precursor derived, Arg2+)

**Figure S4: Myeloid compartment fosters basal cell growth in vitro and in vivo** (A) Addition of various cytokines to basal cell cultures (n=6) and quantification of sphere size after 7-days in culture. (B) Confirmation of IL4R$\alpha$ deletion after CMV-Cre-GFP addition to IL4R$\alpha^{flox/flox}$ basal cell cultures n=3; (D) Analysis of CIAE-NOG animals in Polidocanol injury timeline and (E) Quantification of suprabasal cells (Krt5$^{-}$) after injury; WT n=3 each; CIAE-NOG: n(Sham)=2, n(1d)=2, n(3d)=3; (F) Graphic representation of Ccr2-RFP transgene; (G) Graphical summary;

Table S1: 15-color myeloid marker panel;

| | CD45 | CD19 | CD64 | CD24 | CD11c | MHCII | Siglec F | Ly6C | F4/80 | CD103 | Ly6G | CD11b |
|---|---|---|---|---|---|---|---|---|---|---|---|---|
| Alveolar Mph | + | -- | med | med | hi | lo | hi | -- | + | -- | -- | med |
| Interstitial Mph | + | -- | var | -- | -- | + | lo | -- | var | -- | -- | hi |
| Ly6C+ Mo/Mph | + | -- | lo | -- | -- | -- | -- | hi | lo | -- | -- | hi |
| Ly6C- Mo/Mph | + | -- | -- | var | -- | -- | -- | -- | var | -- | -- | hi |
| CD11b+ DC | + | -- | var | med | -- | + | -- | -- | var | -- | -- | hi |
| CD103+ DC | + | -- | -- | hi | hi | hi | -- | -- | -- | hi | -- | med |
| Neutrophils | + | -- | -- | med | -- | -- | -- | + | -- | -- | + | hi |
| Eosinophils | + | -- | -- | med | -- | -- | med | -- | -- | -- | -- | hi |
| B cells | + | + | -- | + | -- | + | -- | -- | -- | -- | -- | -- |

Table S4: Gene Sets for Gene Enrichement analyses

| Gene Signatures | | | | |
|---|---|---|---|---|
| **S-Phase** | **G2M-Phase** | **Macrophages** | **Monocytes** | **Cluster 7 (IAMs)** |
| Mcm5 | Mki67 | Thbs1 | Lyve1 | Gm10116 |
| Pcna | Tmpo | Fn1 | Selenop | Cxcl16 |
| Tyms | Cenpf | Chil3 | Ccl8 | Tgfbr1 |
| Fen1 | Tacc3 | Vcan | Folr2 | Scgb1a |
| Mcm2 | Fam64a | Ier3 | F13a1 | H2-Eb1 |
| Mcm4 | Smc4 | Vegfa | C1qc | Ms4a7 |
| Rrm1 | Ccnb2 | Osm | C1qa | Cd74 |
| Ung | Ckap2l | Lgals3 | Pf4 | Rpl10-p3 |
| Gins2 | Ckap2 | Ptgs2 | Apoe | H2-Aa |
| Mcm6 | Aurkb | Mmp19 | Cbr2 | Hexb |
| Cdca7 | Bub1 | Clec4n | Gas6 | Gpr65 |
| Dtl | Kif11 | Ctsl | C1qb | Scimp |
| Prim1 | Anp32e | Arg1 | Cd163 | H2-Ab1 |
| Uhrf1 | Tubb4b | Fabp5 | Timp2 | Fyb |
| Mlf1ip | Gtse1 | Cxcl2 | Mrc1 | Glipr1 |
| Hells | Kif20b | Cxcl3 | Stab1 | Cx3cr1 |
| Rfc2 | Hjurp | Spp1 | Maf | Zmynd15 |
| Rpa2 | Cdca3 | Plac8 | Fxyd2 | Gm8730 |
| Nasp | Hn1 | Ccl2 | Cfh | Uba52 |
| Rad51ap1 | Cdc20 | Cxcl1 | Cd209f | Krr1 |
| Gmnn | Ttk | | | Lair1 |
| Wdr76 | Cdc25c | | | Tmem119 |
| Slbp | Kif2c | | | Ppfia4 |
| Ccne2 | Rangap1 | | | Lst1 |
| Ubr7 | Ncapd2 | | | Slamf9 |
| Pold3 | Dlgap5 | | | Gm10020 |
| Msh2 | Cdca2 | | | Tyrobp |
| Atad2 | Cdca8 | | | C1qb |
| Rad51 | Ect2 | | | C1qa |
| Rrm2 | Kif23 | | | Cd72 |
| Cdc45 | Hmmr | | | |
| Cdc6 | Aurka | | | |
| Exo1 | Psrc1 | | | |
| Tipin | Anln | | | |
| Dscc1 | Lbr | | | |
| Blm | Ckap5 | | | |
| Casp8ap2 | Cenpe | | | |
| Usp1 | Ctcf | | | |
| Clspn | Nek2 | | | |
| Pola1 | G2E3 | | | |
| Chaf1b | Gas2l | | | |
| Brip1 | Cbx5 | | | |
| E2f8 | Cenpa | | | |
| | Top2a | | | |
| | Tpx2 | | | |
| | Birc2a5 | | | |
| | Ube2c | | | |
| | Nusap1 | | | |
| | Cdk1 | | | |
| | Hmgb2 | | | |
| | Cks1b | | | |
| | Nuf2 | | | |
| | Cks2 | | | |
| | Ndc80 | | | |
